# SMTrackR: a R/Bioconductor package for mapping protein binding at individual DNA molecules

**DOI:** 10.1101/2025.09.09.674732

**Authors:** Aashna Bansal, Himani Barmola, Shivam Yadav, Satyanarayan Rao

**Affiliations:** Department of Biosciences and Bioengineering, Indian Institute of Technology Roorkee, Uttarakhand 247667, India

**Keywords:** Single Molecule Footprinting, dSMF, protein-DNA binding, TF-nucleosome dynamics, Nanopore Sequencing

## Abstract

Single-molecule assays like NOMe-seq, dSMF and Nanopore are superior to DNase-seq and ATAC-seq as they do not nibble down DNA. Thus, they enable quantification of all three i.e., protein-free, Transcription Factor-bound and histone-complex-bound states. But a user-friendly tool to visualize and quantify such states is lacking. Here, we present, SMTrackR, a Bioconductor package to visualize protein-DNA binding states on individual sequenced DNA molecules. SMTrackR queries the single-molecule footprint database we built and hosted at Galaxy Server. It comprises of BigBed files generated from NOMe-seq, dSMF, and Nanopore (SMAC-seq) datasets. SMTrackR exploits UCSC REST API to query a BigBed file and plot footprint heatmap categorized in different binding states as well as report their occupancies. Additionally, this package generates a Gviz-enabled script to visualize these single-molecules on gene tracks.

**Availability and implementation:** The SMTrackR tool is implemented the statistical programming language R and is available at the GitHub developer site, https://github.com/satyanarayan-rao/SMTrackR. The tool is also available in a web version https://smtrackr.iitr.ac.in. A function is provided to use local BigBed file for users who wish to use unpublished data. A fully automated pipeline to generate such BigBed files is available at https://github.com/satyanarayan-rao/SMF_for_SMThub and https://github.com/satyanarayan-rao/dSMF_for_SMThub.

## 1 Introduction

Pioneering work by (Kelly et al. 2012) enabled mapping of protein binding, predominantly Transcription Factors (TFs) and histone complexes as footprints at individual sequenced DNA molecules. In contrast to nuclease- or transposase-based methods, which fail to capture protein-free state of DNA, these chemical probing-based methods inform all three possible states i.e., protein-free DNA, TF-bound, and histone-bound, at a given locus *in vivo*. Since then, several research groups adopted the technology and even assayed at single-cell level to address key biological questions (Wang et al. 2021). On the same principle, variants like dSMF, SMAC-seq, and Fiber-seq where different methyltransferases whether alone or in tandem were used to achieve about base pair footprint size resolution (Krebs et al. 2017; Stergachis et al. 2020; Shipony et al. 2020; Altemose et al. 2022; Abdulhay et al. 2020; Nanda et al. 2024). Full use of the data, however, has not been utilized to address many more questions or at least as a useful resource. It is potentially due to lack of appropriate database and an interface. Here, we developed a stand-alone front-end, SMTrackR, a R/Bioconductor package to visualize and quantify protein-DNA binding states at a locus of interest from publicly available NOMe-seq, and dSMF datasets. We also provide a web-server version accessible at https://smtrackr.iitr.ac.in. Our approach primarily differs from existing tools in two aspects i) keeping database separate from the front-end and ii) use of UCSC REST API (Kleinendorst et al. 2021; Requena et al. 2019; Rao and Ramachandran 2022). Also, researchers are free to use publicly accessible BigBed tracks as per their need or use their unpublished data to generate desired visualizations. Codes to generate BigBed tracks are available as a snakemake pipeline.

## 2 Materials and Methods

### 2.1 SMTHub

We performed massive sequence analysis on NOMe-seq and dSMF data (∼1TB) from several studies to call protein-DNA binding as footprints. To reliably map all three states, particularly nucleosomes, Illumina sequenced reads ≥ 150 bases was used as a filter (**Figure 1A**; **Supplementary Table 1**). A complete description to annotate footprints on individual sequenced DNA molecules can be found here (Rao and Ramachandran 2022). We made minor changes to adapt the NOMe-seq data with this protocol. Biological replicates were merged to generate high-coverage footprint dataset, and in case of single-cell data, pseudo-bulk footprint datasets were generated by merging corresponding annotated cells, for example all individual 4Cell stage scNOMe-seq bed files with footprint information were merged to generate one 4Cell BigBed track (Wang et al. 2021). When using pseudo-bulk data, we recommend users to turn on the ‘remove_dup’ flag (**Figure S1**).

**Figure 1.**
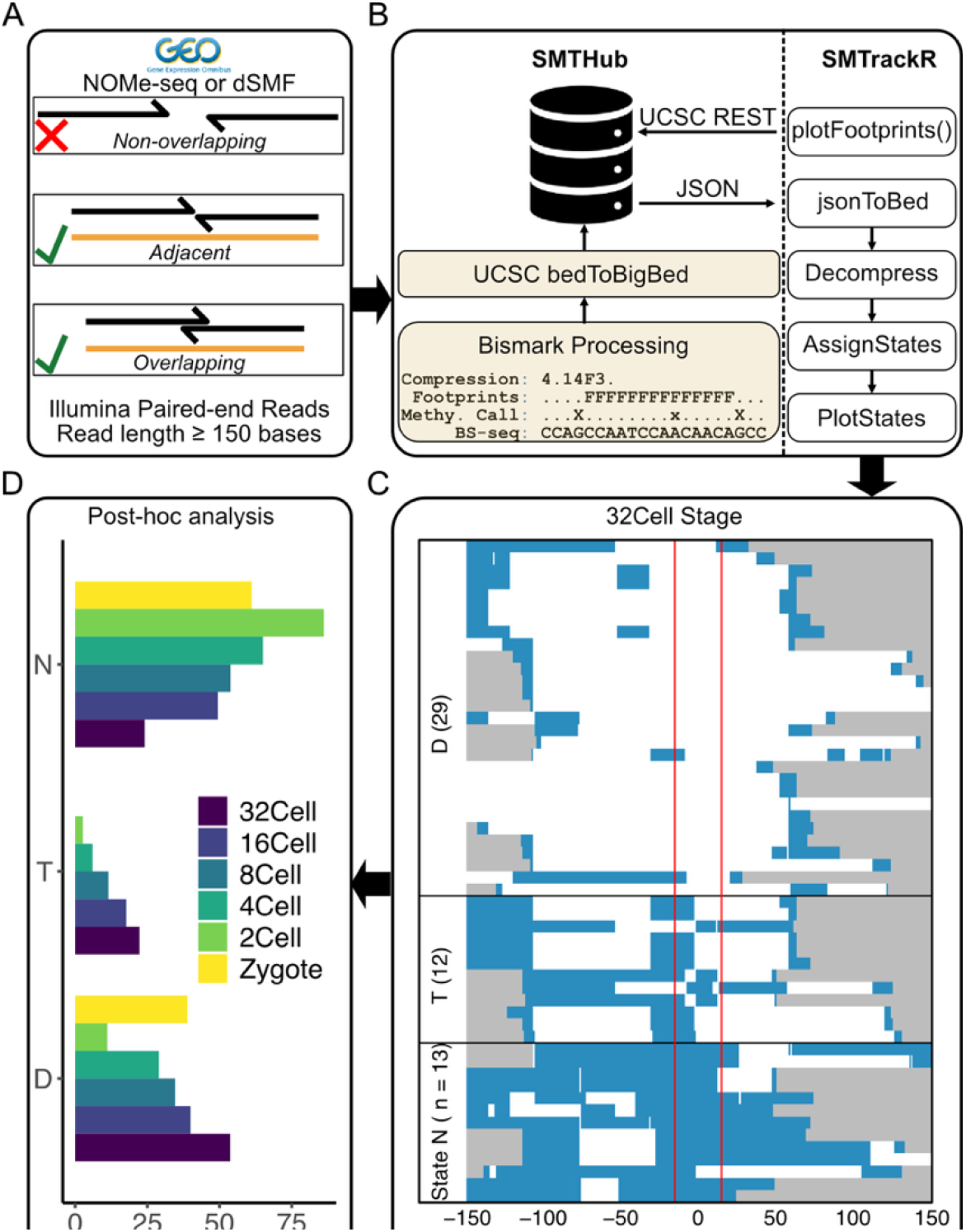
Single-molecule footprinting data processing and chromatin state analysis. **(A)** Sequencing data selection criteria: NOMe-seq or dSMF paired-end reads with at least 150 bp length are selected. Adjacent and overlapping reads are kept for single-molecule footprinting while discarding the non-overlapping reads. **(B)** Bismark is used to call methylation on individual molecules, followed by footprint calls then converted to BigBed and stored in SMTHub. Footprints are retrieved using SMTrackR via the UCSC REST API for visualization and analysis. **(C)** Heatmap generated using SMTrackR for the theoretically derived footprint in the 32-cell stage. Molecules are grouped into three states: *D* (naked DNA), *T* (TF-bound), and *N* (nucleosome-bound), based on methylation patterns. **(D)** Analysis of dynamic changes in occupancy of the proximal region of ZGA genes using SMTrackR across different developmental stages. Nucleosome depletion region (NDR) increases with progression from zygote to the 32-cell stage.

### 2.2 SMTrackR

SMTrackR package queries BigBed files hosted at Galaxy using USCS REST API to fetch footprint information overlapping the locus of interest. It uniquely maps BigBed file with other relevant information like organism of interest, cell-type, conditions etc. The retrieved JSON file is processed which mainly involves: i) expanding the compressed version of footprint information, and ii) assigning binding states based on footprint length and other criteria (Rao and Ramachandran 2022). It then uses base R functions to plot the heatmap. For a better application of the package in visualization context, we have also provided a function ‘generateGvizCodeforSMF’ that writes a Gviz-compatible R script file which places heatmap below the ideogram, axis and genetracks (**Figure S4**) (Hahne and Ivanek 2016). User can change the zoom levels and other parameters as they wish.

### 2.3 Sub-genome suitable for TF footprinting with NOMe-seq

Because NOMe-seq can only utilize GpCs as probes, it has often been used to report primarily nucleosome occupancies. Interestingly, shortest *in silico* footprints, i.e., distance between every first and third cytosines in the GpC context, in mammalian genome sequence alone (tested on mouse and human), we find that about 35 percent (37,679,878 of 107,792,112) are less than 45 base pairs long, and overlaps with regulatory element characterized by ChromHMM (**Figure S2, S3**). Thus, NOMe-seq can reliably map TF-bound states at those loci, particularly for TFs with GpC in their binding motifs. We have compiled a set of TFs (**Supplementary Table 2**) with such motifs and provided *in silico* footprint bed files (for human and mouse genomes) with three contiguous GpCs. A general recommendation is to consider a good number of molecules mapped to region of interest.

## 3 Conclusion

Signals from external stimuli ultimately manifest in action by TFs, activating or repressing a plethora of genes. Such scenarios lead to blocking or unblocking TF binding sites – thus, learning TF-Nucleosome dynamics is key to understanding gene regulation. SMTrackR enables this by mapping all binding states at single-molecule resolution *in vivo*. It covers deeply sequenced dSMF data from *Drosophila melanogaster* S2 cells, a widely used model cell-line for numerous studies. Currently, it supports Illumina-based sequencing datasets and Oxford Nanopore Technology-based methylation calls. In future, we plan to incorporate PacBio long-reads.

## Acknowledgments

Authors thank PARAM Ganga Super Computing Facility Team at IIT Roorkee for their unwavering support for large-volume data processing. Authors deeply acknowledge Georgi K. Marinov, and Anshul Kundaje for sharing SMAC-seq processed methylation calls on Nanopore data from *Saccharomyces cerevisiae*. Authors acknowledge crucial feedback from Vijay Ramani (UCSF), and Ranjith Padinhateeri (IITB).

## Author contribution

AB, HB and SR conceived the project. AB generated all SMTHub datasets. HB analyzed Nanopore data. AB, HB and SR wrote the manuscript. AB, HB, and SR developed the Bioconductor package. SY developed the web-version.

## Funding

AB recognizes MHRD graduate scholarship funding. HB recognizes UCG NET scholarship funding. SR recognizes FIG IITR (FIG-101018) and ANRF (ANRF/ECRG/2024/004316/LS) funding.

## Notes

https://github.com/satyanarayan-rao/SMTrackR

https://github.com/satyanarayan-rao/dSMF_for_SMThub

https://github.com/satyanarayan-rao/SMF_for_SMThub

https://github.com/satyanarayan-rao/GpC_Enriched_Motifs

### Competing Interest Statement

The authors have declared no competing interest.

## References

Abdulhay, Nour J, Colin P McNally, Laura J Hsieh, et al. 2020. “Massively Multiplex Single-Molecule Oligonucleosome Footprinting.” eLife 9 (December): e59404. 10.7554/eLife.59404.

Altemose, Nicolas, Annie Maslan, Owen K. Smith, et al. 2022. “DiMeLo-Seq: A Long-Read, Single-Molecule Method for Mapping Protein–DNA Interactions Genome Wide.” Nature Methods 19 (6): 711–23. 10.1038/s41592-022-01475-6.

Hahne, Florian, and Robert Ivanek. 2016. “Visualizing Genomic Data Using Gviz and Bioconductor.” Methods in Molecular Biology (Clifton, N.J.) 1418: 335–51. 10.1007/978-1-4939-3578-9_16.

Kelly, Theresa K., Yaping Liu, Fides D. Lay, Gangning Liang, Benjamin P. Berman, and Peter A. Jones. 2012. “Genome-Wide Mapping of Nucleosome Positioning and DNA Methylation within Individual DNA Molecules.” Genome Research 22 (12): 2497–506. 10.1101/gr.143008.112.

Kleinendorst, Rozemarijn W. D., Guido Barzaghi, Mike L. Smith, Judith B. Zaugg, and Arnaud R. Krebs. 2021. “Genome-Wide Quantification of Transcription Factor Binding at Single-DNA-Molecule Resolution Using Methyl-Transferase Footprinting.” Nature Protocols 16 (12): 5673–706. 10.1038/s41596-021-00630-1.

Krebs, Arnaud R., Dilek Imanci, Leslie Hoerner, Dimos Gaidatzis, Lukas Burger, and Dirk Schübeler. 2017. “Genome-Wide Single-Molecule Footprinting Reveals High RNA Polymerase II Turnover at Paused Promoters.” Molecular Cell 67 (3): 411–422.e4. 10.1016/j.molcel.2017.06.027.

Nanda, Arjun S., Ke Wu, Iryna Irkliyenko, et al. 2024. “Direct Transposition of Native DNA for Sensitive Multimodal Single-Molecule Sequencing.” Nature Genetics 56 (6): 1300–1309. 10.1038/s41588-024-01748-0.

Rao, Satyanarayan, and Srinivas Ramachandran. 2022. “A Computational Pipeline to Visualize DNA-Protein Binding States Using dSMF Data.” STAR Protocols 3 (2): 101299. 10.1016/j.xpro.2022.101299.

Requena, Francisco, Helena G. Asenjo, Guillermo Barturen, Jordi Martorell-Marugán, Pedro Carmona-Sáez, and David Landeira. 2019. “NOMePlot: Analysis of DNA Methylation and Nucleosome Occupancy at the Single Molecule.” Scientific Reports 9 (1): 8140. 10.1038/s41598-019-44597-2.

Shipony, Zohar, Georgi K. Marinov, Matthew P. Swaffer, et al. 2020. “Long-Range Single-Molecule Mapping of Chromatin Accessibility in Eukaryotes.” Nature Methods 17 (3): 3. 10.1038/s41592-019-0730-2.

Stergachis, Andrew B., Brian M. Debo, Eric Haugen, L. Stirling Churchman, and John A. Stamatoyannopoulos. 2020. “Single-Molecule Regulatory Architectures Captured by Chromatin Fiber Sequencing.” Science 368 (6498): 1449–54. 10.1126/science.aaz1646.

Wang, Yang, Peng Yuan, Zhiqiang Yan, et al. 2021. “Single-Cell Multiomics Sequencing Reveals the Functional Regulatory Landscape of Early Embryos.” Nature Communications 12 (1): 1. 10.1038/s41467-021-21409-8.

