## Supplementary File 1 for "SMTrackR: a R/Bioconductor package for mapping protein binding at individual DNA molecules"

**Supplementary Materials**

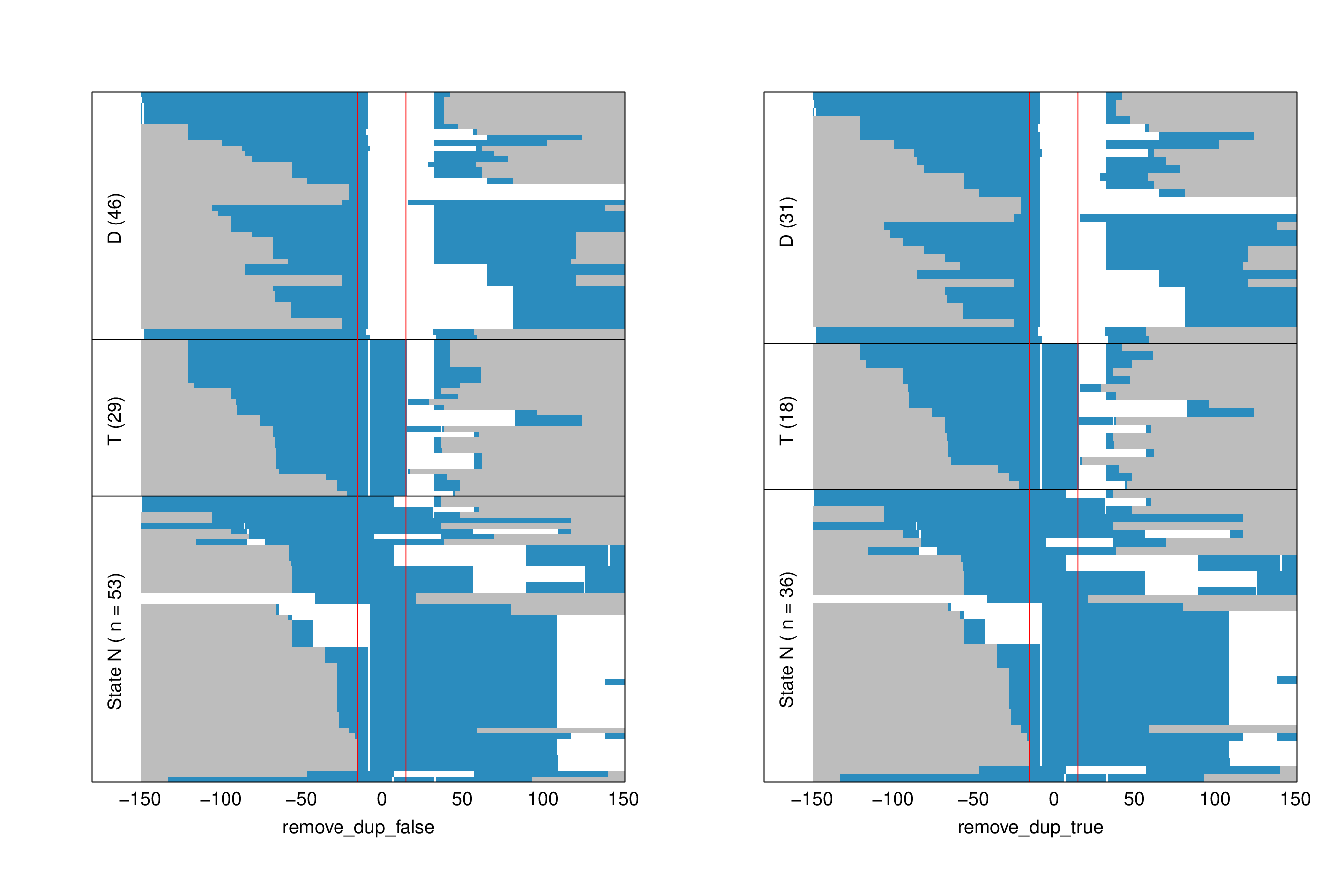

**Figure S1:** **Effect of PCR duplicates in molecule counts.** Panel in the left generate with command: plotFootprints(organism = "mmusculus", model = "8cell", condition = "WT", genome_assembly = "mm10", type = "SMF", chromosome = "chr5", start = "113847750", stop = "113847780", tr = "8cell", label = "remove_dup_true", fp_cap = 50, remove_dup = F), and with remove_dup = T in the right.

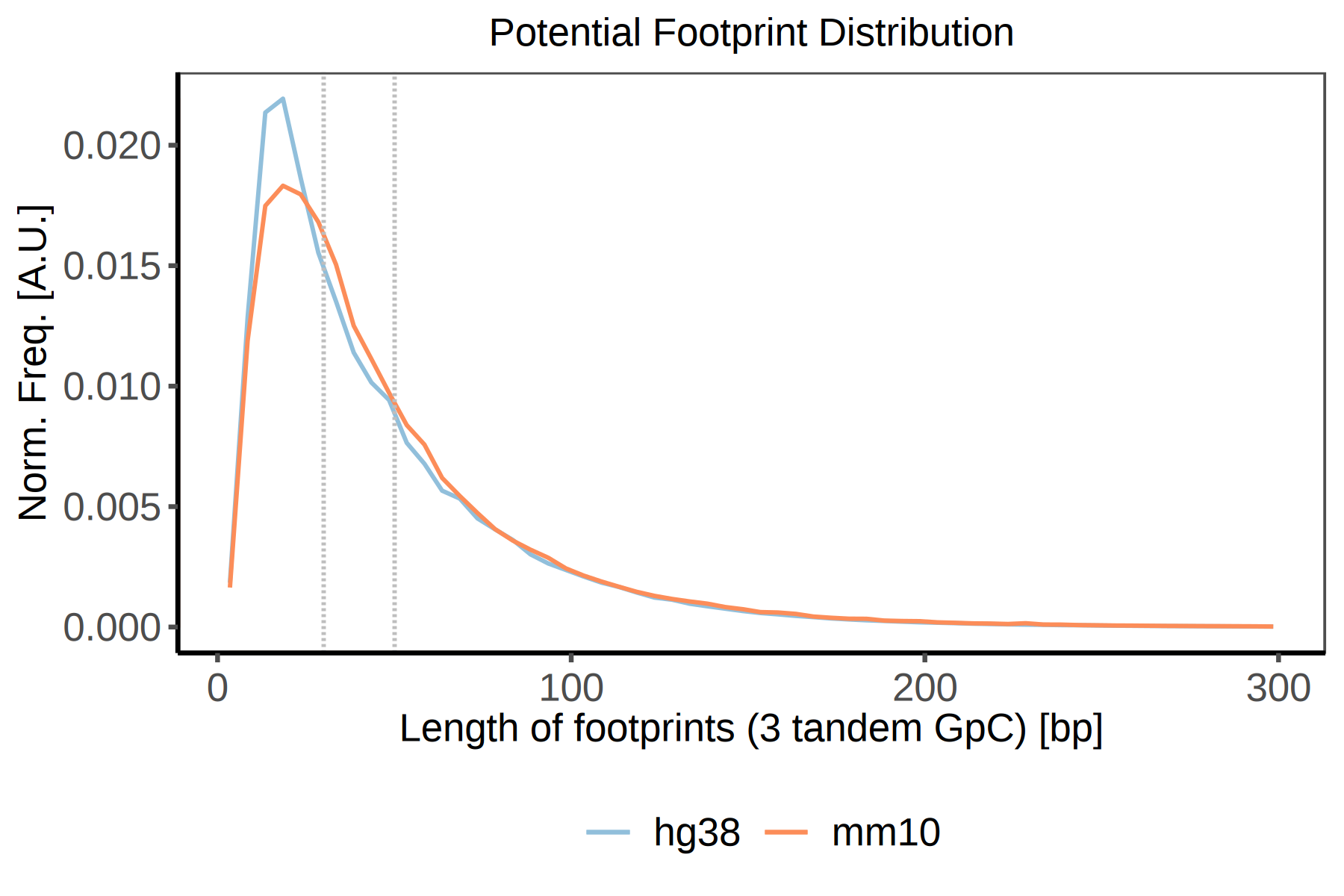

**Figure S2: Theoretical shortest footprint length distribution with GpC as probe**. In Human and Mouse, genome-wide footprints were defined using three contiguous GpCs, potentially giving shortest possible footprint. Vertical grey lines are at 30 and 50 base pairs (bp) respectively.

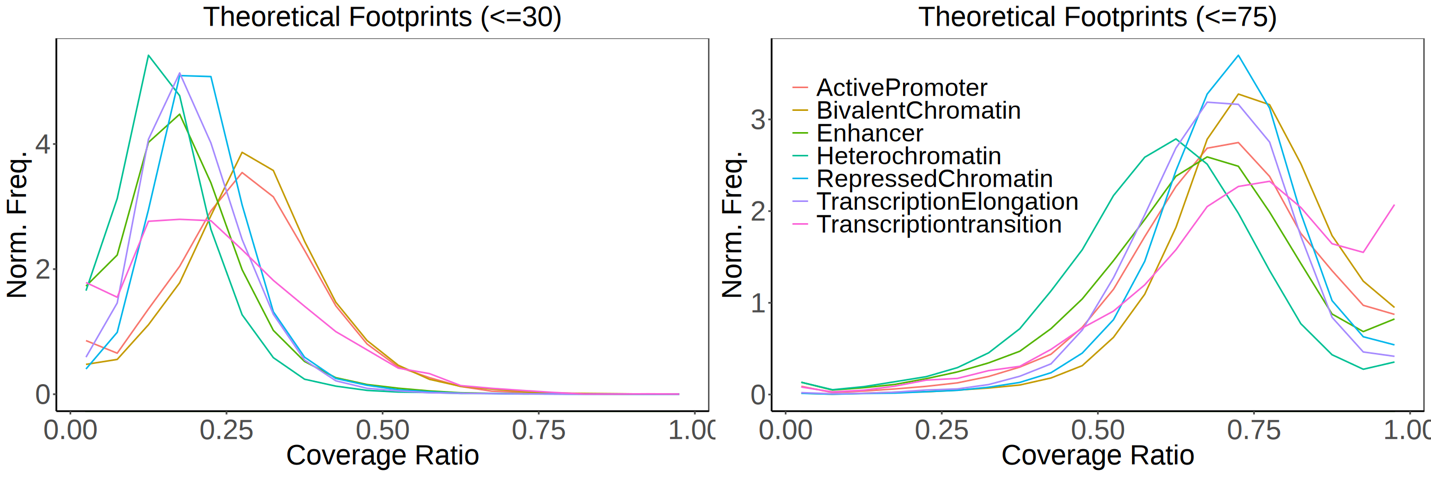

**Figure S3: Shortest theoretical footprint significantly overlaps with chromHMM-defined states.**

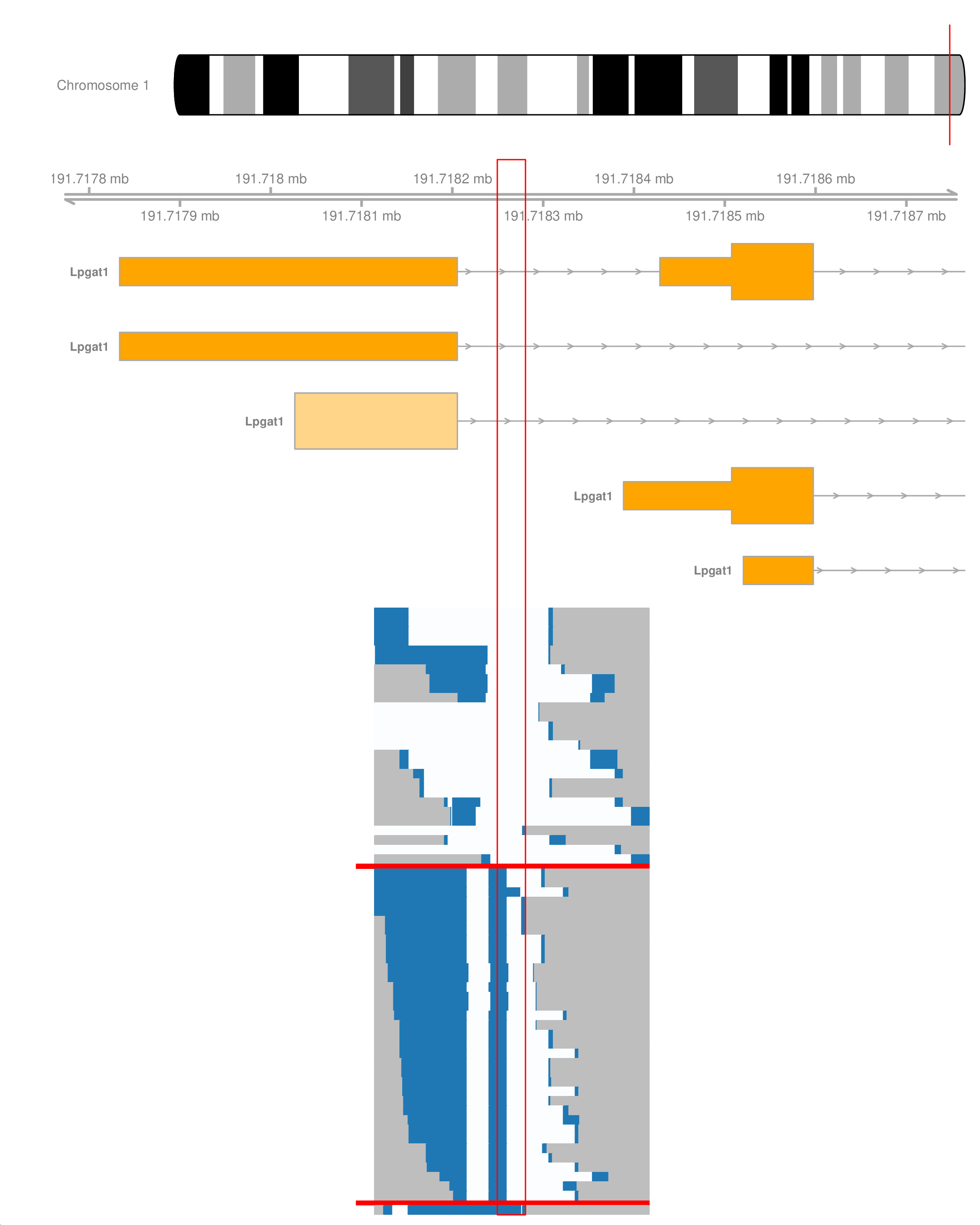

**Figure S4: A plot generated using Gviz compatible code.**
